# Targeting the methyltransferase SETD8 impairs tumor cell survival and overcomes drug resistance independently of p53 status in multiple myeloma

**DOI:** 10.1101/776930

**Authors:** Laurie Herviou, Fanny Izard, Ouissem Karmous-Gadacha, Claire Gourzones, Celine Bellanger, Eva Desmedt, Anqi Ma, Laure Vincent, Guillaume Cartron, Karin Vanderkerken, Jian Jin, Elke De Bruyne, Charlotte Grimaud, Eric Julien, Jérôme Moreaux

## Abstract

Multiple myeloma (MM) is a malignancy of plasma cells that largely remains incurable. The search for new therapeutic targets is therefore essential. Here we show that a higher expression of the lysine methyltransferase SETD8, which is responsible for histone H4K20 mono-methylation, is an adverse prognosis factor associated with a poor outcome in two cohorts of newly diagnosed patients. Remarkably, primary malignant plasma cells are particularly addicted to SETD8 activity. Indeed, pharmacological inhibition of this enzyme by the chemical compound UNC0379 demonstrated a significantly higher toxicity in MM cells compared to normal cells from the bone marrow microenvironment. Moreover, RNA sequencing and functional studies revealed that SETD8 inhibition induces a mature non-proliferating plasma cell signature and an activation of the p53 canonical pathway, which together leads to an impairment of myeloma cell proliferation and survival. However, UNC0379 treatment triggers a deadly level of replicative stress in p53 deficient MM cells, indicating that the cytotoxicity associated with SETD8 inhibition is independent of the p53 status. Consistent with this, the combination of UNC0379 with the conventional cytotoxic agent melphalan strongly enhances DNA damage and overcomes drug resistance in myeloma cells. Thus, targeting SETD8 could be of therapeutic interest to improve MM treatment in high-risk patients independently of the p53 status.

## INTRODUCTION

Multiple Myeloma (MM) is a cancer of terminally differentiated plasma cells characterized by bone destruction, renal failure, and anemia^1^. It is the second most common hematological malignancy after non-Hodgkin lymphoma^2^. Over the past 15 years, several advances in treatments have led to a significantly higher survival of intensively treated patients. This involves a large panel of therapeutic agents, including immunomodulatory drugs and proteasome inhibitors, in combination with autologous stem cell transplantation, alkylating agents (melphalan) and glucocorticoids^3^. Unfortunately, despite the recent progress in MM treatments, most patients will ultimately relapse and develop drug resistance. This cancer thus remains incurable for the majority of patients.

In addition to a wide panel of genetic mutations, recent studies have pinpointed that epigenetic alterations, including aberrant DNA and histone methylation, might be also important players in multiple myeloma development^4^. These recent findings could illuminate new mechanisms central to the genesis of multiple myeloma and offer the possibility to reveal novel approaches and targets for effective therapeutic intervention. For example, the inhibition of the histone H3K27 methyltransferase EZH2 has recently emerged as a potential strategy for the treatment of myeloma^5^. Moreover, classic epigenetic modulating agents, such as histone decatylase inhibitors and DNA methyltransferase inhibitors, are already tested in as monotherapy or in combination with conventional anti-MM agents^6,7^. However, the occurrence of important side effects and the appearance of resistance to these drugs increase the need to identify novel epigenetic targets in multiple myeloma and evaluate their pre-clinical perspective.

SETD8 (also known as SET8, PR-Set7, KMT5A) has been identified as the epigenetic enzyme responsible for the mono-methylation of histone H4 at lysine 20 (H4K20me1) ^8^. SETD8 and H4K20me1 are naturally increased during mitosis and play a critical role in chromatin compaction, gene regulation and cell-cycle progression^8–1010^. In addition, SETD8 could induce the methylation of non-histone proteins, such as the replication factor PCNA and the tumor suppressor p53 ^11,12^. While SETD8-mediated methylation of p53 inhibits apoptosis^12^, PCNA methylation by SETD8 might enhance the interaction with the Flap endonuclease FEN1 and promote cell proliferation^11^. Consistent with this, the overexpression of SETD8 has been reported in many different solid tumors^14–17^ and pharmacological inhibition of SETD8 is sufficient to activate the p53 pro-apoptotic program in neuroblastoma cell lines^18^. This has suggested that this enzyme could be an attractive target to rescue p53 functions in cancers displaying a low incidence of *p53* genetic alterations, as it is the case at early stages in multiple myeloma^19^. However, the role of SETD8 and its incidence in the development of multiple myeloma or any hematological malignancies is not known.

Here, we provide evidence that malignant plasma cells are addicted to SETD8 expression, which is associated with a poor outcome independently of changes in the steady state level of histone H4K20me1. Although inducing p53 canonical pathway, we show that the pharmacological inhibition of SETD8 by the chemical compound UNC0379 also triggers cell-cycle defects and apoptosis in myeloma cells deficient for p53. Finally, the combination of UNC0379 with the cytotoxic agent melphalan strongly enhances DNA damage and overcomes drug resistance, suggesting that targeting SETD8 activity could be beneficial to improve multiple myeloma treatment in high-risk patients independently of their mutational status for p53.

## RESULTS

### SETD8 up-regulation in myeloma is associated with a poor outcome

In order to identify epigenetic factors potentially involved in Multiple Myeloma, we used public affymetrix microarrays gene expression data sets to identify which genes encoding histone-modifying enzymes are differentially expressed between normal bone marrow plasma cells (BMPCs, n=22), purified primary MM cells from newly diagnosed patients (MMCs, n = 345) and human myeloma cell lines (HMCLs, N=42)^19^. As shown in Figure 1A, a significant higher levels of mRNAs encoding the histone H4K20 mono-methyltransferase SETD8 was found in HMCLs compared to BMPCs and MMCS (Fig 1A). Furthermore, although we did not observe a statistical difference with BMPCs, *SETD8* mRNA levels appeared heterogeneous in MMCs (Figure 1A) ranging from 65 to 63338 in affymetrix signal. This contrasted *SETD8* mRNA levels was not restrained to a particular MM molecular sub-type (**supplementary Figure S1**). Consistent with these observations, immunoblot analysis with specific SETD8 antibody showed that HMCLs and some MMCs displayed higher SETD8 protein levels compared with non-cancerous plasma cells (PC) (Figure 1B). These higher SETD8 protein levels were independent to the natural cell-cycle fluctuation of the enzyme^9^, as the levels of the mitotic marker histone H3-S10 phosphorylation were relatively similar in all tested samples (Figure 1B). SETD8 is known as the unique enzyme responsible for the mono-methylation of histone H4 at lysine 20 (H4K20me1)^8^. Yet, the levels of H4K20me1 remained roughly unchanged in HMCLs and MMCs displaying higher SETD8 levels (Figure 1B), indicating that SETD8 up-regulation does not trigger an increase in the steady state level of histone H4K20me1 in multiple myeloma.

**Figure 1.**
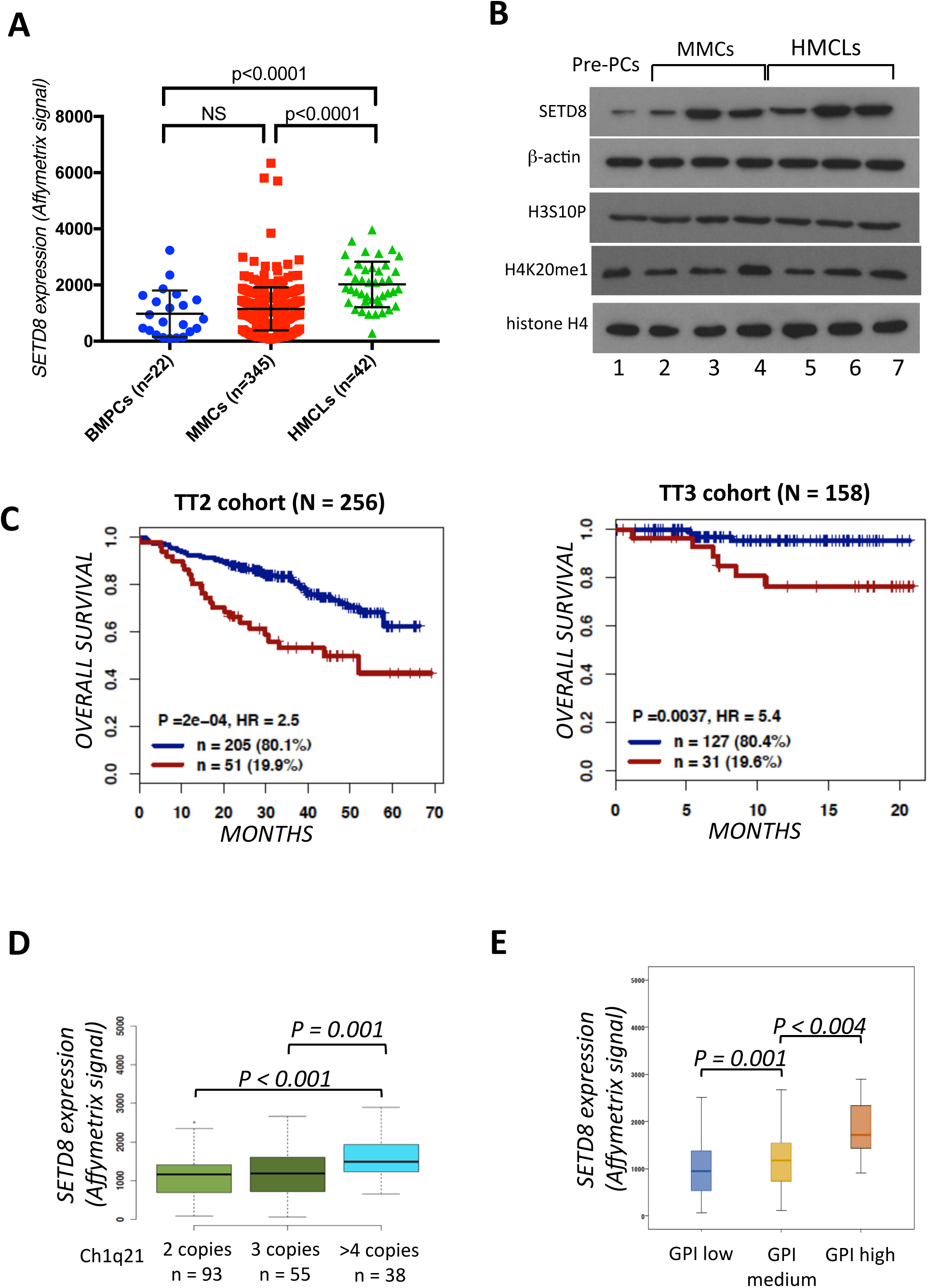
SETD8 expression is a prognosis value in MM. **(A)** SETD8 gene expression in BMPCs, patients’ MMCs and HMCLs. Data are MAS5-normalized Affymetrix signals (U133 plus 2.0 microarrays). Statistical difference was assayed using a student t-test. **(B)** Immunoblot analysis of indicated proteins in total lysates from normal pre-plasmablasts (pre-PC), MMCs and HMCLs. β-actin and histone H4 were used as loading controls. **(C)** Overall survival and Event Free Survival of newly diagnosed MM patients (UAMS-TT2, N=345; and UAMS-TT3, N=158) who’s MMCs highly expressed SETD8 gene. The splitting of the patients into two groups according to SETD8 expression in MMCs was done using the Maxstat algorithm. **(D)** *SETD8* expression in 186 patients of the UAMS-TT2 cohort showing Ch1q21 copy number aberration. **(E)** *SETD8* expression in MMCs (patients at diagnosis) presenting low, medium or high gene expression-based proliferation index (GPI).

We next investigated the prognostic value of SETD8 up-regulation in two independent cohorts of previously untreated MM patients (UAMS-TT2 n=256 and UAMS-TT3 n=158 cohorts), using Maxstat R algorithm. High *SETD8* mRNA levels were predictive of both shorter event free survival and overall survival in the two cohorts (Figure 1C). Consistent with these results, *SETD8* mRNA levels were significantly increased in patients harboring Chr1q21 gain (Figure 1D) or presenting a high gene expression-based proliferation index (GPI) (Figure 1E), molecular features being associated with a poor outcome in MM patients. However, *SETD8* expression did not correlate with MMC plasma cell labeling index (PCLI) in a cohort of 101 newly diagnosed patients (**supplementary Figure S2)**, suggesting that SETD8 up-regulation is likely not a hallmark of MMC proliferation status. Gene set expression analysis (GSEA) of patients with high *SETD8* expression highlighted a significant enrichment of genes involved in IRF4 targets, MYC-MAX targets, MAPK pathway and DNA repair (**Supplementary Figure S3**, P<0.001), suggesting that SETD8 up-regulation correlates with changes in signaling pathways involved in MM pathophysiology. Furthermore, SETD8 expression is significantly higher in MM cell of patients at relapse compared to newly diagnosed patients, underlining a potential role of SETD8 in drug resistance (**Supplementary Figure S4**). Altogether, these data reveal that SETD8 is overexpressed in myeloma and this up-regulation is associated with a poor outcome and deregulation of major signaling pathways in MM patients.

### UNC0379-mediated SETD8 inhibition leads to cell-cycle defects and apoptosis in MMCs

To determine the biological significance of SETD8 up-regulation in MM pathophysiology, the effects of the small-molecule SETD8 inhibitor UNC0379 were examined in eight different HMCLs representative of the disease. UNC0379 is a well-characterized substrate-competitive inhibitor selective for SETD8^15,18,22^. As shown in Figure 2A, UNC0379 treatment was sufficient to inhibit the growth of all HMCLs in a dose dependent manner with an average half maximal inhibitory concentration (IC_50_) ranging from 1.25 to 6.3 μM (Figure 2A). To determine the molecular mechanisms of this HMCL growth inhibition, the effects of UNC0379 treatment on SETD8 activity, cell proliferation and survival were examined using immunoblot and flow cytometry assays in XG7 and XG25 MM cell lines, which displayed similar IC_50_ values for UNC0379. As shown in Figure 2B, analysis of whole cell extracts after 24 hours of treatment showed a strong decrease in H4K20me1, but not of SETD8 and histone H4, thereby demonstrating the rapid and efficient inhibition of SETD8 activity (Figure 2B). In the following hours, this SETD8 inhibition was associated with cell-cycle defects, as shown by an accumulation in G1 phase and a decrease in DNA replication (S) phase 48 hours after treatment (Figure 2C). At later time points, these cell cycle defects was followed by the activation of apoptosis, as measured by the appearance of 33% and 26% of annexin-V positive XG7 and XG25 cells (Figure 2D).

**Figure 2.**
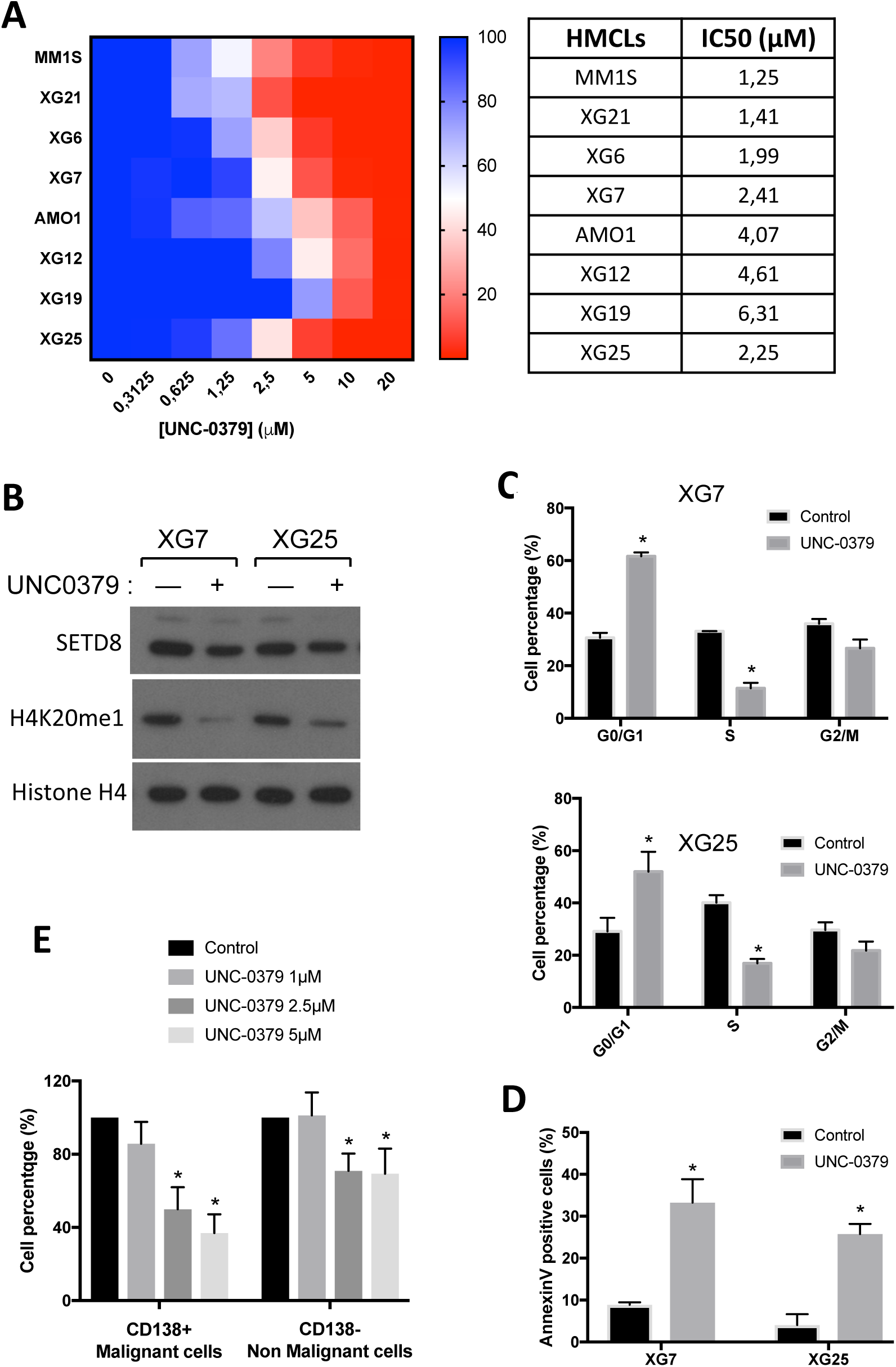
SETD8 inhibitor UNC0379 is highly toxic in malignant plasma cells. **(A)** Graphical representation of HMCLs viability upon exposure with various concentration of UNC-0379. Color scale represents cell viability, from the highest (blue) to the lowest (red) values. Value of IC50 of each HMCL tested is indicated. Data are mean values ± standard deviation (SD) of five experiments determined on sextuplet culture wells. **(B)** Immunoblot analysis of SETD8, Histone H4 and H4K20me1 protein levels in XG7 and XG25 cells untreated or treated with 3 μM of UNC0379 for 24 hours. **(C)** Quantitation of cell-cycle distribution of control (untreated) and UNC0379-treated XG7 and XG25 HMCLs 48 hours after treatment. After short-pulse of BrdU incorporation, cell-cycle was analyzed by FACS using DAPI and anti-BrdU antibody. Results are representative of three independent experiments. (*) indicates a significant difference compared to control cells using a Wilcoxon test for pairs (P ≤ 0.05). (**D**) Quantitation of apoptosis in control and UNC-0379-treated XG7 and XG25 HMCLs by flow cytometry with AnnexinV-PE staining and 96h after UNC0379 treatment. Data shown are mean values ± SD of 4 separate experiments. Statistical analysis was done with a paired t-test. (*) indicates a significant difference compared to control cells using a Wilcoxon test for pairs (P ≤ 0.05). **(E)** Percentage of *in vitro* cultivated primary MM cells (CD138+ tumor cells) and bone marrow microenvironment (CD138-non-malignant cells) upon increasing concentration of UNC0379 treatment for 4 days. Data shown are mean values of 8 patient samples. (*) indicates a significant difference compared to control cells using a Wilcoxon test for pairs (P ≤ 0.05).

To confirm that SETD8 is essential for growth and viability of malignant plasma cells, bone marrow (BM) from MM patient samples were cultured with recombinant interleukin 6 in presence or not of UNC0379. Five days after treatment, the percentage of myeloma and non-myeloma cells was then measured by flow cytometry after staining with anti-CD138 antibody that specifically recognizes plasma cells. As shown in Figure 2E, the median number of CD138 positive malignant plasma cells were decreased in a dose-dependent manner, with 50% and 63% of reduction with 2.5 μM and 5 μM of UNC0379 respectively (*P* = 0.001 and *P* < 0.0001; N=8), whereas non-tumoral BM cells were less sensitive with a decrease of 26% and 24% at these UNC0379 concentrations. In addition, to support the feasibility of preclinical studies with SETD8 inhibitors, primary 5T33vv murine MM models^23^ were treated with growing concentrations of UNC0379 for 24h hours before cell viability analysis. As observed with human primary MM cells (Figure 2E), UNC0379 treatment results in significant reduction of 5T33vv viability in a dose dependent manner (**Supplementary Figure S5**). We therefore conclude that malignant plasma cells are particularly addicted to SETD8 activity and that the pharmacological inhibition of this epigenetic enzyme is highly toxic, leading rapidly to cell growth inhibition and apoptosis.

### SETD8 inhibition impairs MM cell proliferation together with activation of p53 target gene pathways

SETD8 has been involved in all nuclear processes that use DNA as a matrix, including a critical role in the regulation of gene expression^8^. In order to gain insights into the mechanisms contributing to SETD8-mediated MM cell growth inhibition and death, we therefore performed transcriptome analysis in XG7 and XG25 HMCLs after SETD8 inhibition using RNA sequencing (RNAseq). To identify gene expression alterations caused by SETD8 inhibition in HMCLs, we isolated total RNAs 18 hours after UNC0379 treatment or 48 hours after shRNA-induced SETD8 silencing, when H4K20me1 decrease did not trigger DNA replication defects yet^24,25^. A common signature of 820 up-regulated genes and 360 down-regulated genes was found in UNC0379-treated and SETD8-depleted cell lines compared to the untreated and shRNA control cell lines (Fold change > 2; FDR ≤ 0.05) (**supplementary figure S6** and **Supplementary Table 1**). Gene Set Enrichment Analysis (GSEA) identified, as the most down-regulated pathways, genes involved in cell-cycle, stem cells, proliferating plasmablasts and MM proliferating molecular subgroup together with genes repressed upon loss of the histone H3K27 methyltransferase EZH2 (Figures 3A and 3B). Conversely and consistent with the presence of a functional p53 in XG7 and XG25 cell lines, a significant positive enrichment was found for p53 target genes, including *p21* and *GADD45A* genes, and genes overexpressed in mature BMPCs versus plasmablasts (Figures 3C and 3D). A positive enrichment for genes normally down-regulated by c-MYC, EZH2, DNA methylation or histone deacetylases (HDAC) was also observed (Figure 3C), which might suggest some impairments of chromatin silencing pathways upon SETD8 inhibition in MM cells. As shown in figure 3E, immunoblot analysis of UNC0379-treated XG7 HMCLs confirmed the increased levels of p53 and p21 proteins upon UNC0379 treatment. However, we did not observe a significant phosphorylation of the histone variant H2A.X (Figure 3E), suggesting p53 activation in XG7 MM cells treated with UNC0379 occurs in absence or low level of DNA damage. Altogether, these results data indicate that the cytotoxic effects of SETD8 inhibition in p53-proficient HMCLs are associated with mature non-proliferating plasma cell transcriptional signature and activation of the p53 canonical pathway.

**Figure 3.**
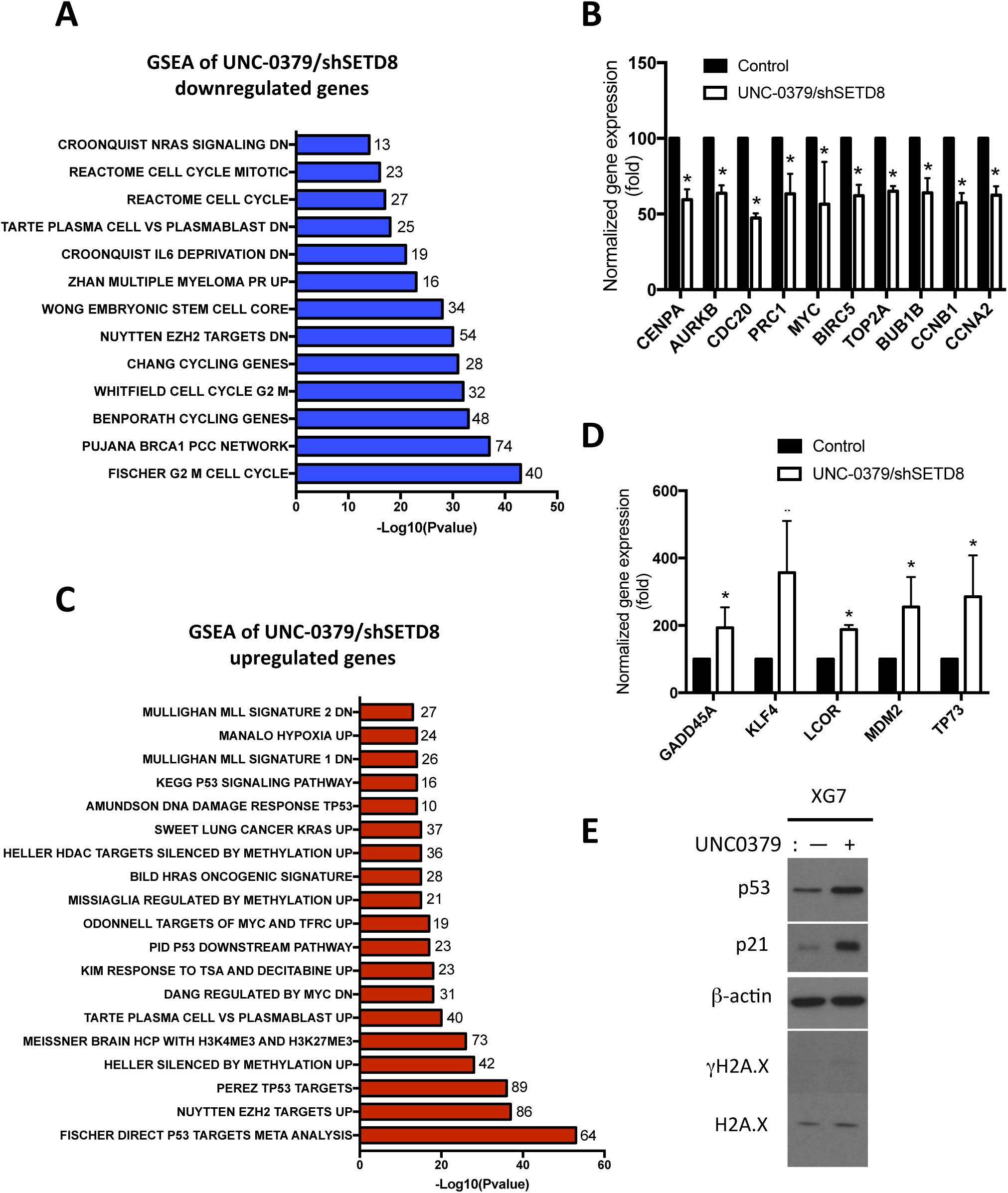
Gene expression changes in HMCLs upon genetic or pharmacological inhibition of SETD8. **(A)** Molecular signatures of UNC-0379 and shSETD8 downregulated genes compared to control was investigated using GSEA Database (all curated gene sets), and relevant pathways are presented (FDR q-value ≤ 0.05). (**B**) Bar-plot representing the fold expression (UNC-0379 condition over control) of genes related to (A) chart pathways. (**C**) Molecular signatures of UNC-0379 and shSETD8 up-regulated genes compared to control was investigated using GSEA Database (all curated gene sets), and relevant pathways are presented (FDR q-value ≤ 0.05) **(D)** Bar-plot representing the fold expression (UNC-0379 condition over control) of genes related to (C) chart pathways. **(E)** Immunoblot analysis of indicated proteins in total lysates from UNC-0379-treated or untreated XG7 HMCL. β-actin and H2A.X were used as loading controls.

### The cytotoxicity of UNC0379 treatment is not dependent on p53 in malignant plasma cells

Previous studies showed that UNC0379-induced cell death depends on the activation of p53 in neuroblastoma cancer models (NB), thereby rending p53-deficient NB cells more resistant to SETD8 inhibition (Veschi et al. 2017). To determine whether the cytotoxic effects of the pharmacological inhibition of SEDT8 mediated by UNC0379 was also dependent on p53 activation in MM cells, we compared the UNC0379 response in p53 wild-type HMCLs (n=7) and p53-deficient HMCLs (n=5)^20,21^ that displayed similar *SETD8* expression levels (Figure 4A). Noted of, there was no significant correlation between *SETD8* and *P53* expression in HMCLs (**Supplementary Figure S7**). Although IC_50_ values for p53-deficient HMCLs following UNC0379 treatment were more heterogeneous, we found no statistical difference with the group of p53 wild-type HMCLs (Figure 4B). We also observed no statistical difference in the IC_50_ values of UNC0379 according to the expression levels p53 in HMCLs, whatever their p53 status (**Supplementary Figure S8**). Altogether, these results suggested that the absence of a functional p53 did not alter the sensitivity of malignant plasma cells to the pharmacological inhibition of SETD8 activity. To verify this hypothesis and gain insights into the mechanisms by which UNC0379 could impair the viability of malignant plasma cells independently of p53, we examined by immunoblot and flow cytometry analysis the cytotoxic effects of UNC0379 treatment on p53-proficient XG7 HMCL transduced with high-titer of lentivirus encoding either a p53 shRNA or an irrelevant control shRNA. Both cell lines showed similar IC_50_ values for UNC0379 (3.7 _μ_M in control shRNA and 4.8 μM in P53 shRNA expressing cells). Immunoblot analysis confirmed the efficient p53 depletion in p53 shRNA XG7 HMCL relative to control cells (Figure 4C). Consistent with results in Figure 3E, UNC0379 treatment led to an up-regulation of p53 without detectable DNA damage in shRNA control XG7 cells (Figure 4C). In contrast, higher levels of DNA damage and replicative stress were observed in UNC0379-treated XG7 cells depleted for p53, as evidenced by the increased levels of both phosphorylated histone variant H2A.X and checkpoint protein CHK1 (Figure 4C) and by an accumulation of cells in G2/M phase of the cell cycle (Figure 4D). This was followed by the appearance of a high percentage of apoptotic cells, similarly to the percentage detected in UNC0379-treated control shRNA XG7 cells displaying p53 activation (Figure 4E). Importantly, a similar DNA damage signature was observed in XG1 HMCL naturally harboring a mutated inactive p53 (**Supplementary Figure S9**). Thus, these results indicated that UNC0379-induced cytotoxicity in MM cells is independent on p53 status and that a high level of SETD8 likely protects MM cells from spontaneous intrinsic DNA damage and replicative stress.

**Figure 4.**
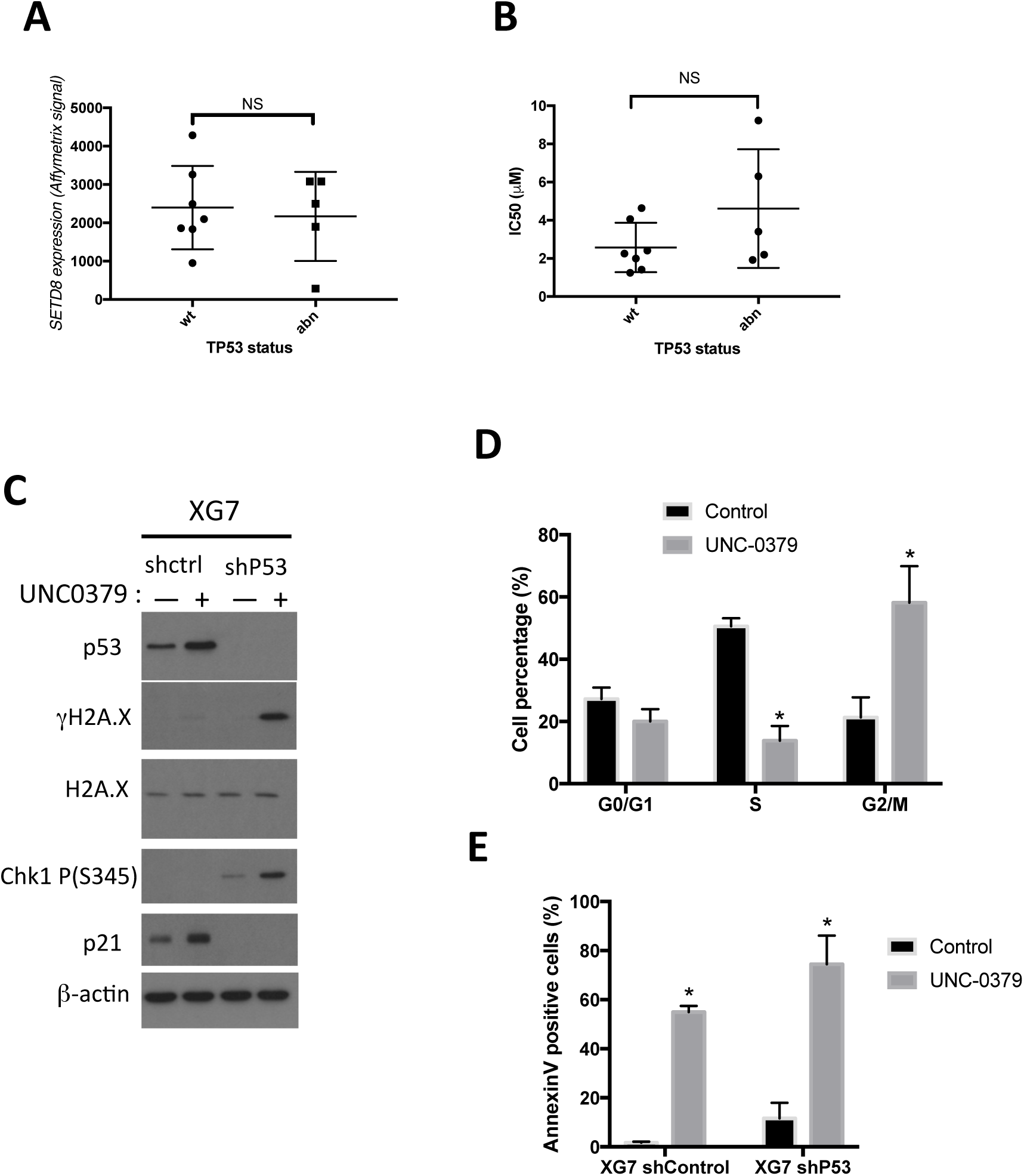
UNC0379 toxicity is independent of P53. **(A)** Comparison of *SETD8* expression according to HMCLs *TP53* status. **(B)** Comparison of UNC-0379 IC_50_ (μM) according to HMCLs *TP53* status. **(C)** Immunoblot analysis of indicated proteins in total lysates from UNC-0379-treated (5μM) or untreated XG7-shControl and XG7-shTP53 HMCLs. β-actin and H2A.X were used as loading controls. (**D**) Cell cycle of control and 48h UNC-0379-treated (5μM) XG7-shTP53 HMCL was analyzed by flow cytometry using DAPI, BrdU incorporation and labelling with an anti-BrdU antibody. Results are representative of three independent experiments. * indicates a significant difference compared to control cells using a Wilcoxon test for pairs (P ≤ 0.05). (**E**) Apoptosis induction in control and UNC-0379-treated (5μM) XG7-shControl and XG7-shTP53 HMCLs was analyzed with AnnexinV-PE staining by flow cytometry after 96h treatment. Data shown are mean values ± SD of 4 separate experiments. Statistical analysis was done with a paired t-test. * indicates a significant difference compared to control cells using a Wilcoxon test for pairs (P ≤ 0.05).

### Pharmacological inhibition of SETD8 synergizes with melphalan

The results presented above indicate that the pharmacological inhibition of SETD8 could constitute a promising strategy to improve multiple myeloma treatment by increasing DNA damage and genomic instability. To further explore this possibility, we investigated whether SETD8 inhibition could enhance the cytotoxicity of melphalan and overcome resistance to this alkylating agent widely used in MM treatment. To this end, we first measured by FACS the levels of apoptotic cells in sensitive (MelphS) and melphalan-resistance (MelphR) XG7 MM cell lines treated with either 5 μM of melphalan, 3 μM of UNC0379 or a combination of the two compounds. Note of, the levels of SETD8 and TP53 expression levels were similar in both cell lines (**Supplementary Figure S10**). Whereas UNC0379 alone slightly increased the percentage of annexinV-positive MelphS and MelphR cells, the combination with Melphalan significantly enhanced this percentage in MelphR cells and, to a lesser extent, in MelphS cells (**Figure 5A**). Remarkably, the overcome resistance to melphalan upon UNC0379 treatment in MelpR cells was associated with an enhancement of DNA breaks, as observed by a higher level of phosphorylated histone variant H2AX and by an increase in the number of DNA damage-induced 53BP1 foci in these cells (**Figures 5B and 5C**). However, although we observed an activation of the cell-cycle checkpoint p21 (**Figure 5C**), UNC0379 sensitized HMCLs to melphalan treatment was independently from the presence of p53 (**Figure 5D**). Altogether, these results indicate that a low dose of UNC0379 is sufficient to significantly increase the percentage of melphalan-induced MM cell death in a p53 dependent manner and to overcome resistance to this alkylating agent, thereby demonstrating the therapeutic interest to target SETD8-mediated lysine methylation in multiple myeloma whatever their p53 status.

**Figure 5.**
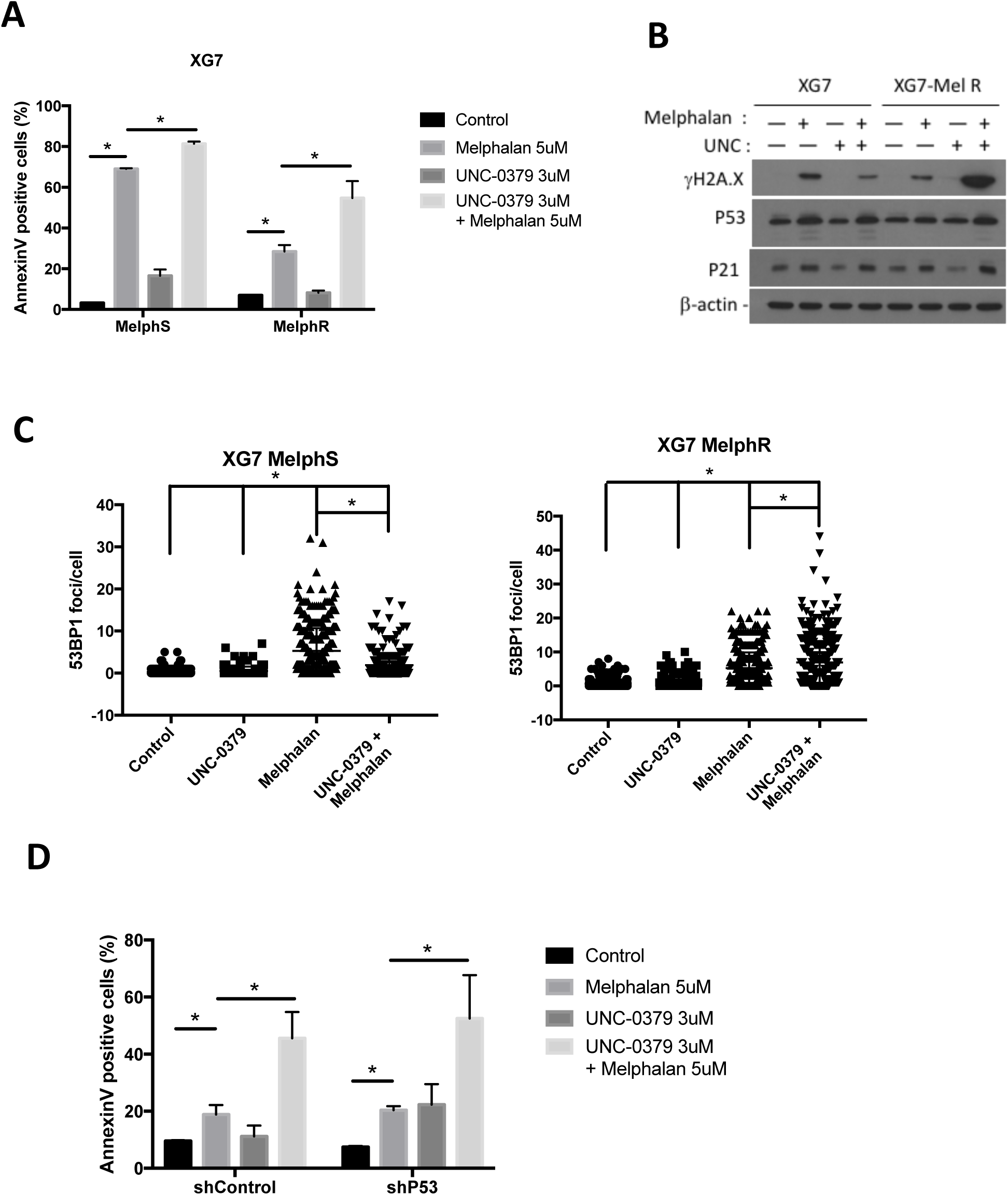
UNC-0379 treatment sensitizes HMCLs to Melphalan-induced DNA damage. **(A)** Apoptosis induction in XG7 HMCLs sensitive (MelphS) or resistant (acquired resistance, MelphR) to Melphalan, after 96h of treatment with Melphalan (5μM), UNC-0379-treated (3μM) or the combination of the two drugs. AnnexinV-PE staining was analyzed by flow cytometry after 96h treatment. **(B)** Immunoblot analysis of indicated proteins in total lysates from XG7 MelphS or MelphR HMCLs, treated with Melphalan (5μM), UNC-0379-treated (3μM) or the combination of the two drugs for 24h. β-actin was used as loading control. **(C)** 53BP1 foci were observed in XG7 MelphS or MelphR HMCLs, treated with Melphalan (5μM), UNC-0379-treated (3μM) or the combination of the two drugs for 24h. Number of foci per cell was quantified using ImageJ software (mean number of cells counted: 300). Statistical significance between conditions was assessed using Student paired t-test (*: pvalue<0.05). **(D)** Apoptosis induction in XG7-shControl and XG7-shTP53 HMCLs after 96h of treatment with Melphalan (5μM), UNC-0379-treated (3μM) or the combination of the two drugs. AnnexinV-PE staining was analyzed by flow cytometry.

## DISCUSSION

This study unveils for the first time a role of the lysine methytransferase SETD8 in multiple myeloma (MM) patho-physiology and drug resistance. We have shown that elevated levels of SETD8 are associated with a poor prognosis in two large cohorts of newly diagnosed patients and correlates with the deregulation of regulatory nodes involved in multiple myeloma, including IRF4, MYC/MAX, MAPK and DNA repair pathways (Figure 1). Moreover, patients displaying higher SETD8 expression are associated with specific molecular features, such as increase copy of Chr1q21 or a high GPI index (Figure 1). Consistent with this, primary MM cells are particularly addicted to SETD8 activity and the recently developed small-molecule inhibitor of SETD8, UNC0379, demonstrated a significantly higher toxicity in MM cells compared to normal cells from the bone marrow microenvironment (Figure 2). From a mechanistic point of view, our RNA-seq results show that the genetic or pharmacological inhibition of SETD8 in MM cell lines results in the activation of a mature non-proliferating plasma cell signature and of the p53 canonical pathway (Figure 3). However, UNC0379-induced cytotoxicity does not necessarily require p53 activation as p53 wild-type and deficient MM cells display similar sensitivity to SETD8 inhibition (Figures 3 and 4). Finally, in order to support future pre-clinical study, we have shown that a low dose of UNC0379 enhances melphalan-induced cell death and overcomes the resistance associated with this DNA-damaging conventional cytotoxic agent in both p53 wild-type and deficient MM cells (Figure 5). Altogether, these results provide evidence for SETD8 inhibition as a potential novel therapeutic strategy in multiple myeloma independently from the mutational p53 status of these tumors.

An up-regulation of SETD8 is not specific to multiple myeloma, as it has also been observed in different types of solid tumors^11^, such as papillary thyroid cancer^16^, breast carcinoma^14,26^, and childhood tumors of the nervous system^15,18^. However, the mechanisms that contribute to elevated levels of SETD8 in cancer still remained unclear. In neuroblastoma, a higher-level of SETD8 could be due in part to DNA copy-number gains at chr12q24, the region that encompasses *SETD8* encoding gene^26^. Another non-exclusive mechanism is related to the deregulation of oncogenic c-MYC pathway, since *SETD8* encoding gene has been found as a transcriptional target of c-MYC^27^ and functionally required to mediate MYC-induced cell growth^15^. It is widely established that c-MYC is a key regulator in MM with deregulations related to translocations, gains and amplification, mutations in RAS genes and MYC transcription or translation activation^28^. Interestingly, while chr12q24 gains are not present in myeloma, we identified here a significant enrichment of c-MYC/MAX target genes in MM patients characterized by high *SETD8* expression, thereby supporting a potential role of this oncogenic pathway in SETD8 deregulation. Furthermore, SETD8 depletion or inhibition results in up-regulation of genes repressed upon c-MYC expression (Figure 3C) suggesting that SETD8 might also participate in deregulation of c-MYC functions in multiple myeloma.

An unexpected result of our study is that the up-regulation of SETD8 in malignant plasma cells was not associated with significant changes in the steady state level of the mono-methylation of histone H4K20 (H4K20me1), the main target of SETD8^8^. This result is independent from cell-cycle progression, since changes in the levels of SETD8 do not correlate with cell proliferation (figure 1B and supplementary figure S2). One hypothesis is that an overexpression of the H4K20me1-specific demethylase PHF8, observed in many cancers including hematopoietic malignancies^29,30^, could attenuate SETD8 activity on histone H4K20^31^. Thus, although we cannot rule out that some local changes in H4K20me1 levels may occur at specific genetic loci, our results indicate that the global level of this epigenetic modification is not a valuable marker of SEDT8 activity and functions in multiple myeloma.

Recent studies have suggested that the role of SETD8 in cancer might involve the methylation of other substrates than histone H4. Thus, Takawa et al. proposed that SEDT8 can methylate the proliferating cell nuclear antigen PCNA and thus favor HeLa cell proliferation^11^; but this result was never further confirmed in other cell types. Additionally, SETD8 can induce the mono-methylation of the tumor suppressor p53 at lysine 382 (p53K382), which attenuates its pro-apoptotic and growth arrest functions^12,18^. Hence, in neuroblastoma, inhibition of SETD8 by UNC0379 leads to cell death in a dependent p53 manner and p53K382 is important for this phenotype^18^. Our RNA-seq and functional studies in MM cells shows that the genetic or pharmacological inhibition of SETD8 leads to the activation of p53 pathway, which correlates with an increased in p53 and p21 protein levels, G1/S arrest followed by cell death. However, in contrast to neuroblastoma^18^, UNC0379-induced cytotoxicity in MM cells is not necessarily dependent on p53 activation and, despite several attempts, we were unable to detect p53K382me1 in MM cells. Few DNA breaks are generally sufficient to activate P53 functions^32,33^. Since the first impact of loss of SETD8 activity is excessive chromatin relaxation and progressive DNA damage after exit of mitosis^34^, we propose that the activation of p53 we detect in MM cells upon SETD8 inhibition is mainly triggered by cellular stresses during G1 progression. This would also explain why UNC0379-mediated SETD8 inhibition is very toxic in P53-deficient MM cells, where the absence of a functional p53-mediated G1 arrest allows damaged cells to pursue into S-phase and accumulate a deadly level of DNA breaks, likely caused by improper replication fork progression^24^. Altogether, our findings are consistent with a model in which UNC0379-mediated cell death is triggered by p53 activation in p53-proficient MM cells and rather associated with high replicative stress in cancer cells deficient for p53, thereby rending multiple myeloma sensitive to SEDT8 inhibition whatever the mutational status of p53.

In spite of effective therapeutic protocols developed in MM, drug resistance remains a major concern. In this regard, we report here that high *SETD8* expression is associated with a poor prognosis in patients treated by high-dose melphalan and autologous stem cell transplantation. SETD8 expression is also significantly up-regulated in patients at relapse compared to newly diagnosed patients. In agreement with this, UNC0379 treatment potentiates the cytotoxicity of melphalan and overcomes melphalan drug resistance in MM cells, underlining the interest to target SETD8 to improve the treatment of MM. Melphalan is an alkylating cytotoxic agent used in MM patients treated by high dose chemotherapy and hematopoietic stem cell autograft and in non-transplantable patients, in combination with other molecules. The protective effect against melphalan provided by SETD8 expression is associated with the poor prognosis of MM patients characterized by elevated levels of SETD8.

Deletion of the short arm of chromosome 17 (del 17p) is associated with a poor outcome in MM independently of treatment regimen^35–39^. Interestingly, MM cell toxicity mediated by SETD8 inhibitor is mainly p53-independent. The frequency of events targeting p53 (del 17p, *TP53* mutation or double hits) increases during the progression of MM and consecutive relapses underlining a selection of cells harboring TP53 abnormalities in association with resistance to treatment. TP53 bi-allelic events are also associated with a dramatic impact on MM patients’ survival after relapse^40^. Therapies inducing significant toxicity of p53 defective MM cells are needed. Our results demonstrate the therapeutic interest of SETD8 inhibitor to target p53 deficient MM cells by increasing replicative stress and DNA damages. Accordingly, combination of SETD8 inhibitor with melphalan could be of clinical interest, notably in newly diagnosed patients presenting del17p and/or *SETD8* overexpression and eligible to high dose melphalan and ASCT. Thus, SETD8 inhibition appears of therapeutic interest to overcome drug resistance and improve the treatment of MM patients at relapse independently of the p53 status.

## MATERIALS AND METHODS

### Primary multiple myeloma cells

Bone marrow samples were collected after patients’ written informed consent in accordance with the Declaration of Helsinki and institutional research board approval from Heidelberg and Montpellier University hospital. Bone marrow were collected from 206 patients treated with high dose Melphalan (HDM) and autologous stem cell transplantation (ASCT) and this cohort is termed “Heidelberg-Montpellier” (HM) cohort. Patients’ MMCs were purified using anti-CD138 MACS microbeads (Miltenyi Biotec, Bergisch Gladbach, Germany) and their gene expression profile (GEP) obtained using Affymetrix U133 plus 2.0 microarrays as described^41,42^. The CEL files and MAS5 files are available in the ArrayExpress public database (E-MTAB-372). The structural chromosomal aberrations, as well as numerical aberrations were assayed by fluorescence in situ hybridization (iFISH). We also used publicly available Affymetrix GEP (Gene Expression Omnibus, accession number GSE2658) of a cohort of 345 purified MMC from previously untreated patients from the University of Arkansas for Medical Sciences (UAMS, Little Rock, AR), termed in the following UAMS-TT2 cohort. These patients were treated with total therapy 2 including HDM and ASCT^43^. We also used Affymetrix data from total therapy 3 cohort (UAMS-TT3; n=158; E-TABM-1138)^44^ of 188 relapsed MM patients subsequently treated with bortezomib (GSE9782) from the study by Mulligan *et* al^45^. The mouse 5T33MMvv cells originated spontaneously in aging C57BL/KaLwRij mice and have since been propagated *in vivo* by intravenous transfer of the diseased marrow in young syngeneic mice as described^23,46^.

### Treatment of primary MM cells

Bone marrow of patients presenting with previously untreated MM (n = 8) at the university hospital of Montpellier was obtained after patients’ written informed consent in accordance with the Declaration of Helsinki and agreement of the Montpellier University Hospital Centre for Biological Resources (DC-2008-417). Mononuclear cells were treated with or without UNC-0379 (1μM, 2.5μM or 5μM) and MMC cytotoxicity were evaluated using anti-CD138-phycoerythrin monoclonal antibody (Immunotech, Marseille, France) as described^47^.

### Human Myeloma Cell Lines (HMCLs)

XG human myeloma cell lines (HMCLs) were cultured in the presence of recombinant IL-6 as previously described^20^. XG7 Melphalan-resistant cells were generated by successively treating the sensitive parental XG7 cells with 0.6μM of Melphalan for 3 months. JJN3 was kindly provided by Dr Van Riet (Brussels, Belgium), JIM3 by Dr MacLennan (Birmingham, UK) and MM1S by Dr S. Rosen (Chicago, USA). AMO-1, LP1, L363, U266, OPM2, and SKMM2 were purchased from DSMZ (Braunsweig, Germany) and RPMI8226 from ATTC (Rockville, MD, USA). HMCLs were authenticated according to their short tandem repeat profiling and their gene expression profiling using Affymetrix U133 plus 2.0 microarrays deposited in the ArrayExpress public database under accession numbers E-TABM-937 and E-TABM-1088^20^.

### Establishment of shRNA expressing HMCLs

Control and p53 shRNA sequences were cloned in the pLenti4-EZ-mIR plasmid as previously described^48^. SETD8 and associated control shRNA sequences were cloned into a puromycin retroviral vector RNAi Ready pSiren as described^24^. Retroviral particles were produced in 293FT cells. Briefly, 293FT cell line was cultured in Dulbecco’s modified Eagle’s medium and supplemented with 10% defined fetal bovine serum, 500μg/ml geneticin, 4mM *L*-glutamine, and 1mM MEM sodium pyruvate. The day before transfection, cells were plated into a 10cm tissue culture plate to 90%-95% confluence. 9μg of ViraPower packaging mix (Invitrogen) and 9μg of lentiviral plasmids were co-transfected into 293FT cells using 36μl Lipofectamine 2000 reagent (Invitrogen). Forty-eight hours later, culture supernatants were collected, concentrated 100 fold by ultracentrifugation (20000 g, 4 hours) and viral titers determined. Corresponding HMCLs were transduced with virus and stable transduced cells were obtained by adding zeocin (10 μg/ml) for pLenti4-EZ-mIR and puromycin (2.5μg/ml) for pSIREN viral particles.

### Growth assays and cell cycle analysis

Cells were cultured for 4 days in 96-well flat-bottom microtiter plates in RPMI 1640 medium, 10% FCS, and 2 ng/ml IL-6 (control medium) in the presence of UNC-0379. Cell growth was evaluated by quantifying intracellular ATP amount with a Cell Titer Glo Luminescent Assay (Promega, Madison, WI) using a Centro LB 960 luminometer (Berthold Technologies, Bad Wildbad, Germany). For cell cycle analysis, cells were cultured in 24-well flat-bottomed microtiter plates at 10^5^ cells per well in RPMI1640–10% FCS or X-VIVO 20 culture medium with or without IL-6 (3ng/mL). The cell cycle was assessed using DAPI staining (Sigma-Aldrich, Saint-Louis, MO, USA) and cells in the S phase using incubation with bromodeoxyuridine (BrdU) for 1 h and labeling with an anti-BrdU antibody (APC BrdU flow kit, BD Biosciences, San Jose, CA, USA) according to the manufacturer’s instructions. Flow cytometry analysis was done on a Fortessa flow cytometer (BD, Mountain View, CA, USA).

### Apoptosis assays

Cells were cultured in 24-well, flat-bottomed microtiter plates at 10^5^ cells per well in RPMI1640–10% FCS or X-VIVO 20 culture medium with or without IL-6 (3ng/mL) and appropriate concentration of chemical drugs. After 4 days of culture, cells were washed twice in PBS and apoptosis was assayed with PE-conjugated Annexin V labeling (BD Biosciences) using a Fortessa flow cytometer (BD) following manufacturer protocols.

### RNA sequencing

HMCLs were cultured for 16 hours with or without 5 μM of UNC-0379, or infected with shcontrol or shRNA SETD8 pSIREN vectors for 48h. RNA samples were collected as previously described. The RNA sequencing (RNA-seq) library preparation was done with 150ng of input RNA using the Illumina TruSeq Stranded mRNA Library Prep Kit. Paired-end RNA-seq were performed with Illumina NextSeq sequencing instrument (Helixio, Clermont-Ferrand, France). RNA-seq read pairs were mapped to the reference human GRCh37 genome using the STAR aligner^49^. All statistical analyses were performed with the statistics software R (version 3.2.3; available from https://www.r-project.org) and R packages developed by BioConductor project (available from https://www.bioconductor.org/)^50^. The expression level of each gene was summarized and normalized using DESeq2 R/Bioconductor package^51^. Differential expression analysis was performed using DESeq2 pipeline. P values were adjusted to control the global FDR across all comparisons with the default option of the DESeq2 package. Genes were considered differentially expressed if they had an adjusted p-value of 0.05 and a fold change of 1.5. Pathway enrichment analyses were performed using online the curated gene set collection on the Gene Set Enrichment Analysis software (http://software.broadinstitute.org/gsea/msigdb/index.jsp)^52,53^.

### Gene expression profiling and statistical analyses

Gene expression data were normalized with the MAS5 algorithm and analyses processed with GenomicScape (http://www.genomicscape.com)^54^ the R.2.10.1 and bioconductor version 2.5 programs^50^. Gene Set Expression Analysis (GSEA) was used to identify genes and pathways differentially expressed between populations. Univariate and multivariate analysis of genes prognostic for patients’ survival was performed using the Cox proportional hazard model. Difference in overall survival between groups of patients was assayed with a log-rank test and survival curves plotted using the Kaplan-Meier method (Maxstat R package)^55^.

### 53BP1 Staining-immunofluorescence microscopy

After deposition on slides using a Cytospin centrifuge, cells were fixed with 4% PFA, permeabilized with 0.5% Triton in PBS and saturated with 5% bovine milk in PBS. The rabbit anti-53BP1 (Novus Biologicals - Littleton, CO, USA – NB100304) antibody was diluted 1/300 in 5% bovine milk in PBS, and deposited on cytospins for 60 minutes at room temperature. Slides were washed twice and anti-rabbit alexa 488-conjugated antibody (diluted 1/500 in 5% bovine milk in PBS) was added for 60 minutes at room temperature. Slides were washed and mounted with Vectashield and 1% DAPI. Images and fluorescence were captured with a ZEISS Axio Imager Z2 microscope (X63 objective) (Oberkochen, Germany), analyzed with Omero (omero.mri.cnrs.fr) server and ImageJ software.

### Immunoblot analysis

Cells washed with phosphate-buffered saline (PBS) were lysed in SDS buffer and boiled at 94°C for 5 minutes. After measuring protein quantity by Bradford, equal amounts of protein were resolved by SDS-PAGE, transferred to a nitrocellulose membrane (Millipore) and probed with one of the following antibodies: mouse anti-Chk1 (1:1000, abcam), rabbit anti-p21 (1:500, Cell signaling), rabbit anti-p53 (1:1000, Cell Signaling), rabbit anti-SETD8 (1:1000, Cell Signaling), mouse anti-β-actin (1:20000, Sigma), rabbit anti-H2A.X and anti-phospho-H2A.X-Ser139 (1:1000, Cell signaling), rabbit anti-H4-K20me1 (1:1000 Cell Signaling), and rabbit anti-Histone H4 (1:1000, Cell Signaling). Membranes were then incubated with the appropriate horseradish peroxidase (HRP)-conjugated secondary antibodies. The immunoreactive bands were detected by chemiluminescence (Pierce).

## Legends of supplementary Figures

**Supplementary Figure S1:**
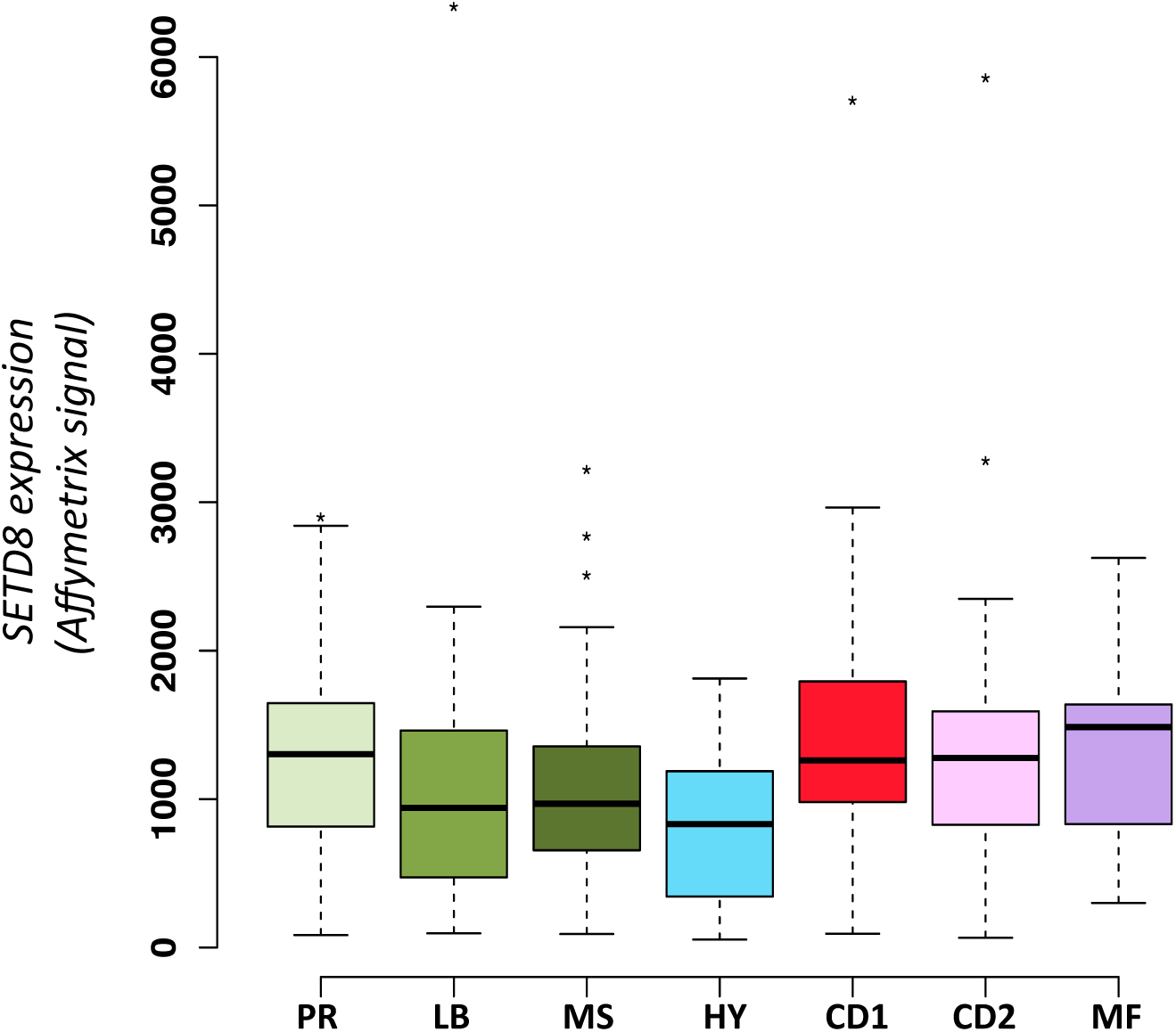
SETD8 expression different subgroup of MM patients. Gene expression profiling of MMCs of the patients of UAMS-TT2 cohort were used. PR: proliferation, LB: low bone disease, MS: MMSET, HY: hyperdiploid, CD1: Cyclin D1-Cyclin D3, CD2: Cyclin D1-Cyclin D3, MF: MAF, MY: myeloid.

**Supplementary Figure S2:**
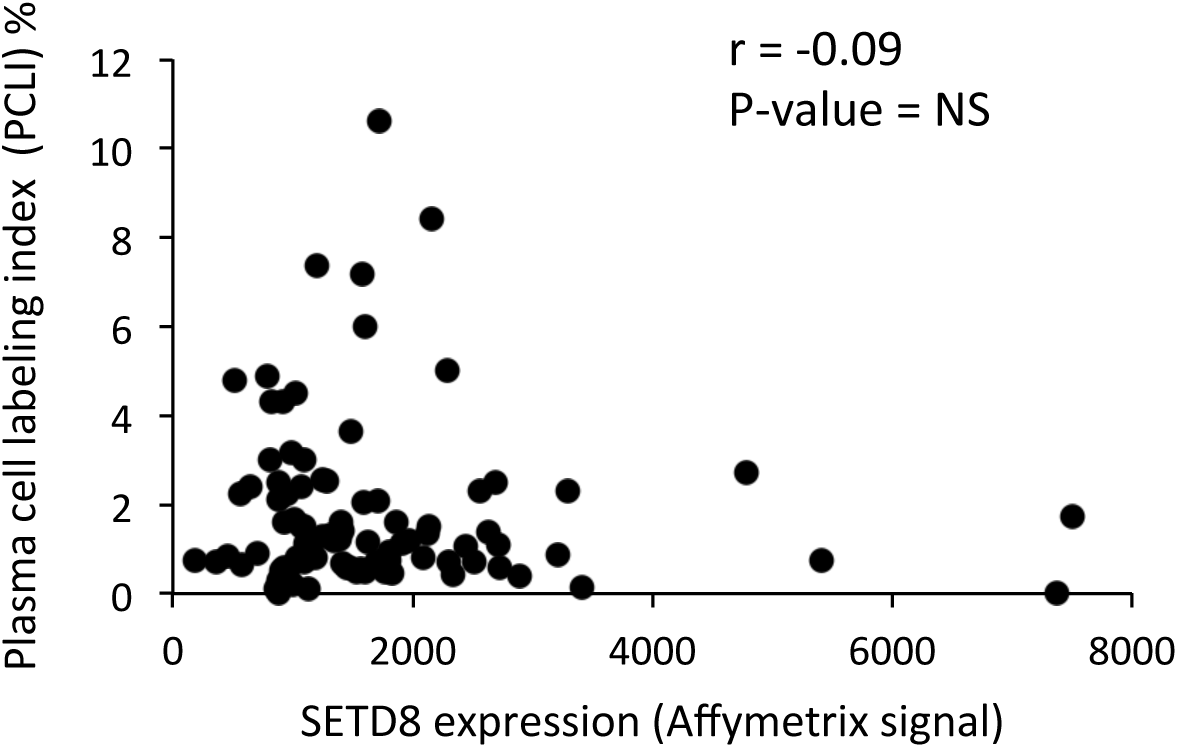
Correlation between SETD8 expression and malignant plasma cell labeling index. Plasma cell labeling index was investigated using Brdu incorporation and flow cytometry in 101 patients at diagnosis.

**Supplementary Figure S3:**
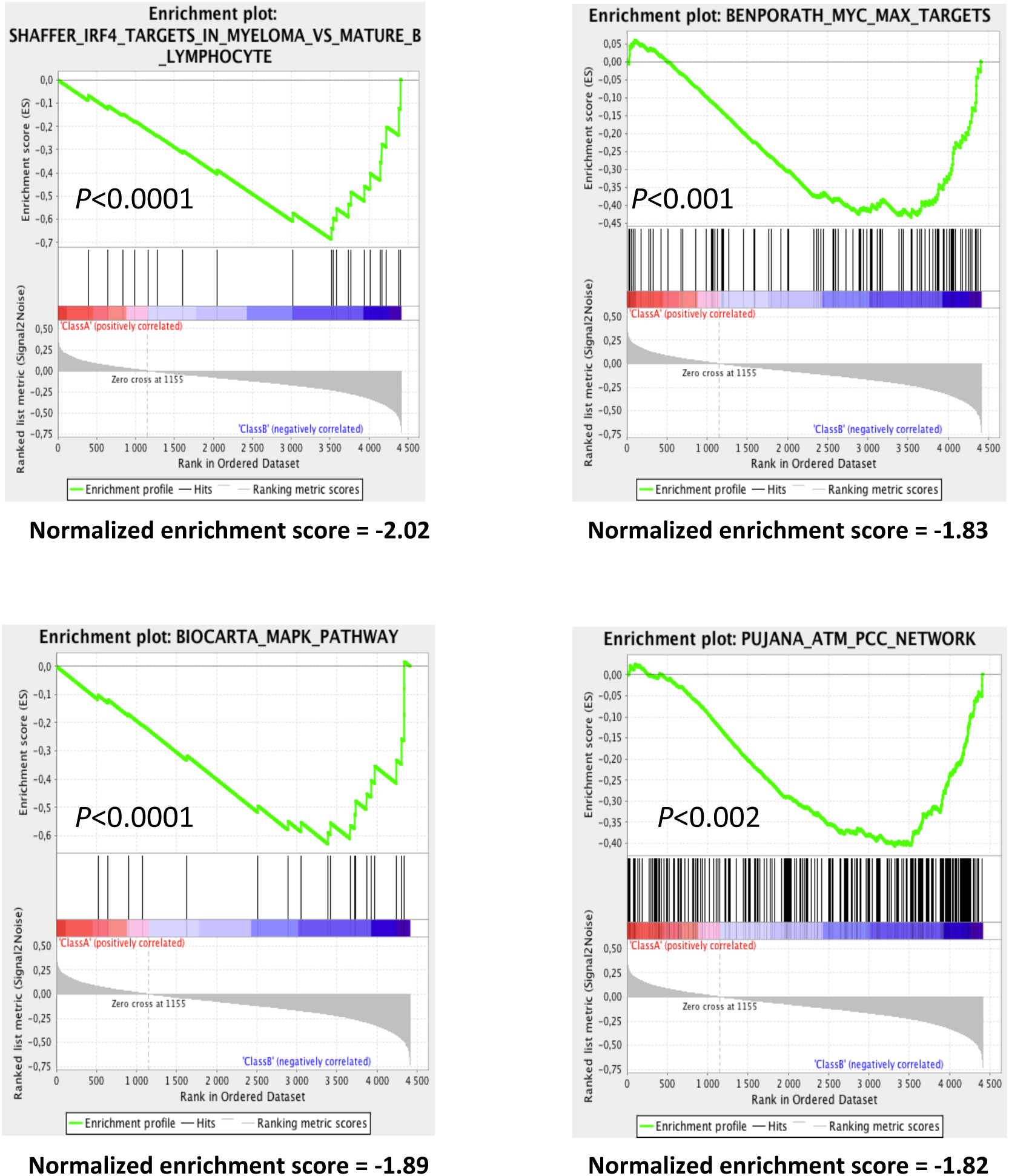
Gene Signature of MM patients with high SETD8 expression. GSEA enrichment plots with the absolute enrichment p value and the normalized enrichment score of the gene set.

**Supplementary Figure S4:**
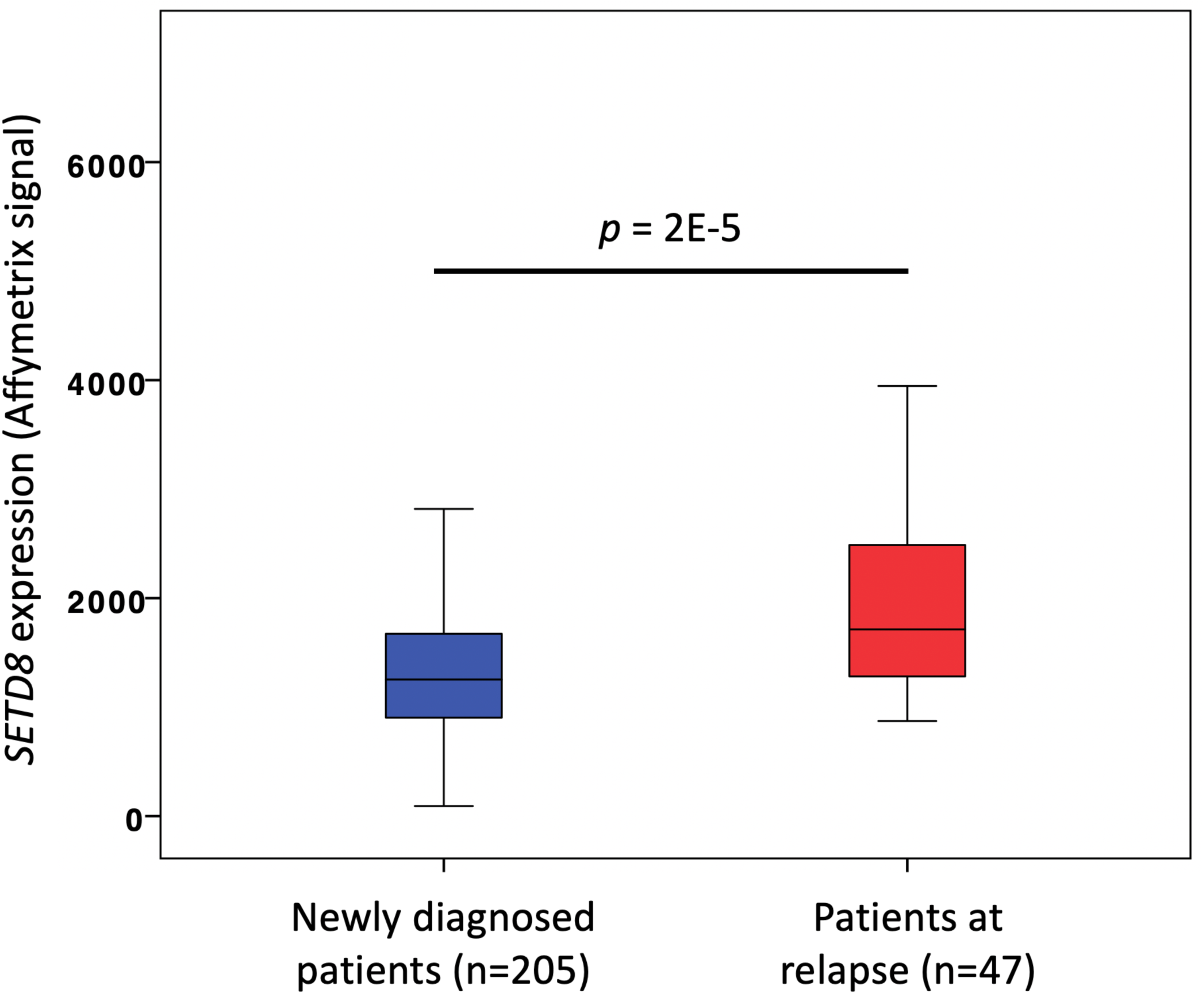
SETD8 expression is significantly higher in MM cells of patients at relapse (n=47) compared to diagnosis (n=205).

**Supplementary Figure S5:**
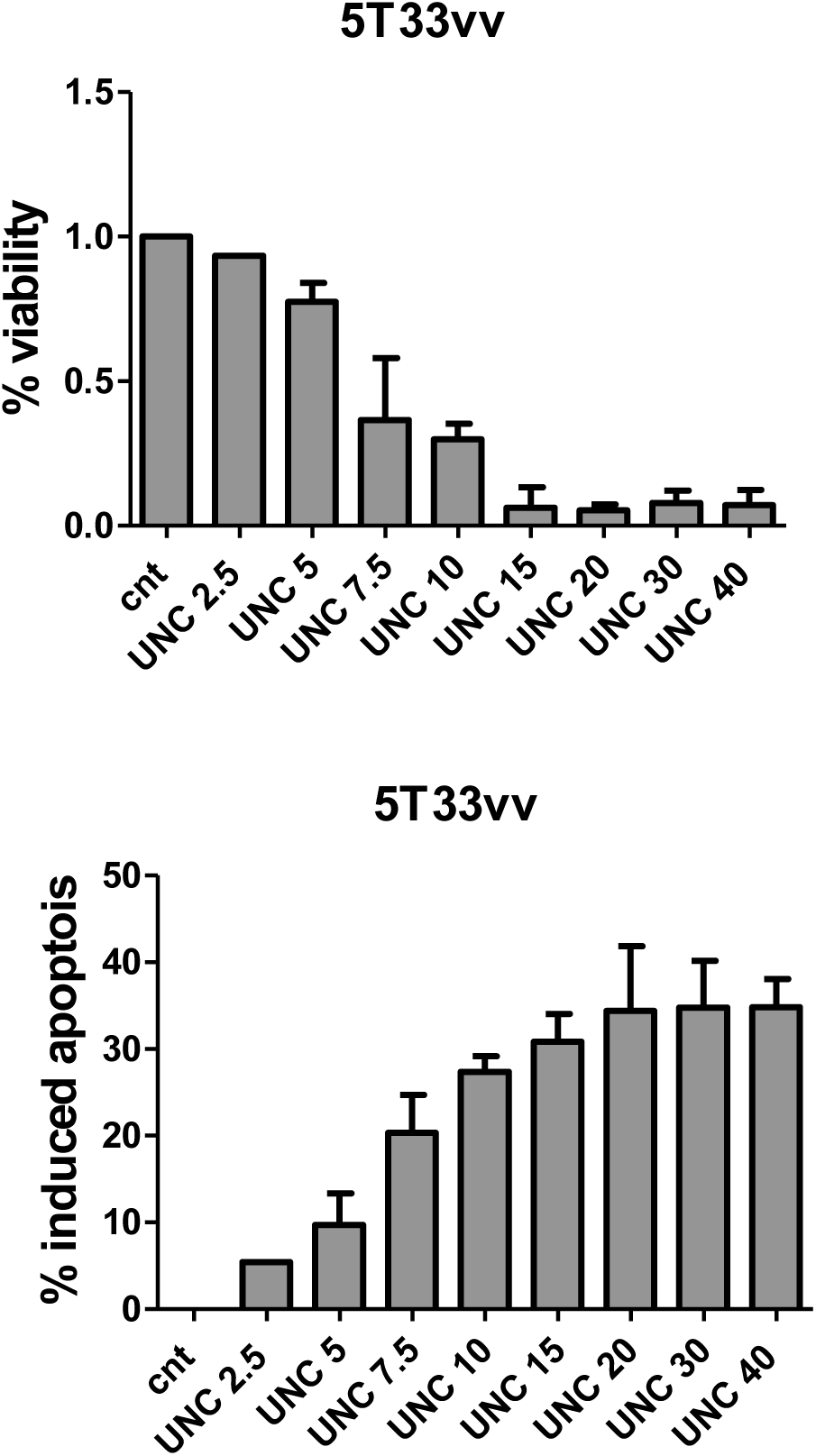
Measure of viability and apoptosis in primary murine 5T3vv cellular models untreated (cnt) or treated with growing concentrations (from 2.5 to 40 μM) of UNC0379 for 24 hours.

**Supplementary Figure S6:**
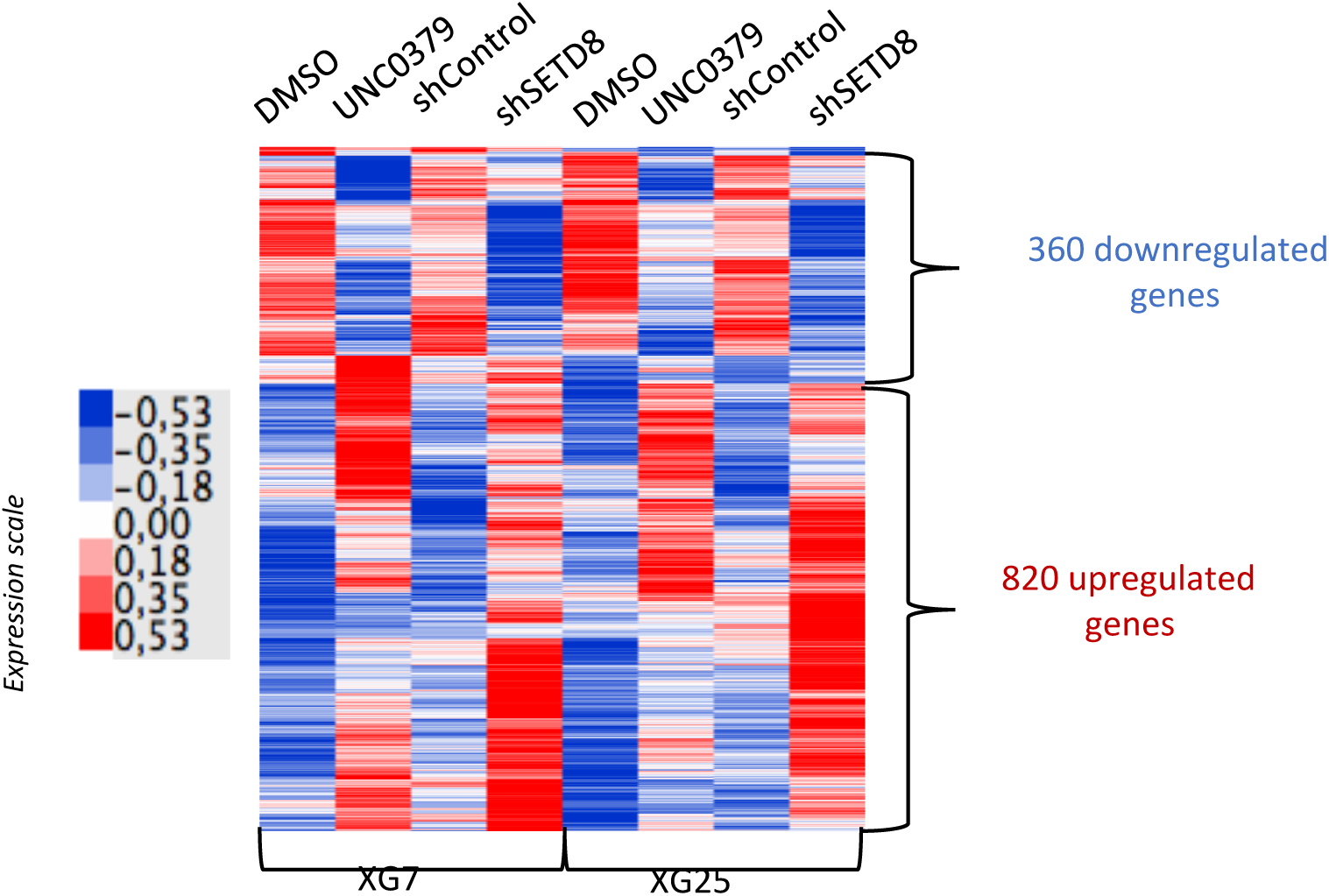
Heatmap of RNA-sequencing rlog expression data. Genes deregulated by UNC-0379 and shSETD8 in XG7 and XG25 HMCLs are represented. Expression scale shows low expression levels in blue and high expression levels in red.

**Supplementary Figure S7:**
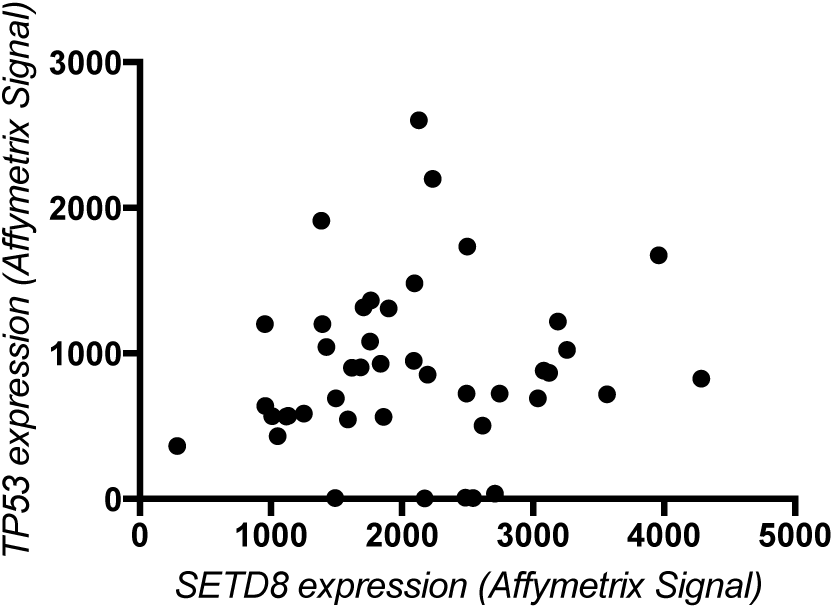
Correlation between *TP53* and *SETD8* expression in HMCLs. *TP53* and *SETD8* expression was obtained from Affymetrix microarrays data previously published.

**Supplementary Figure S8:**
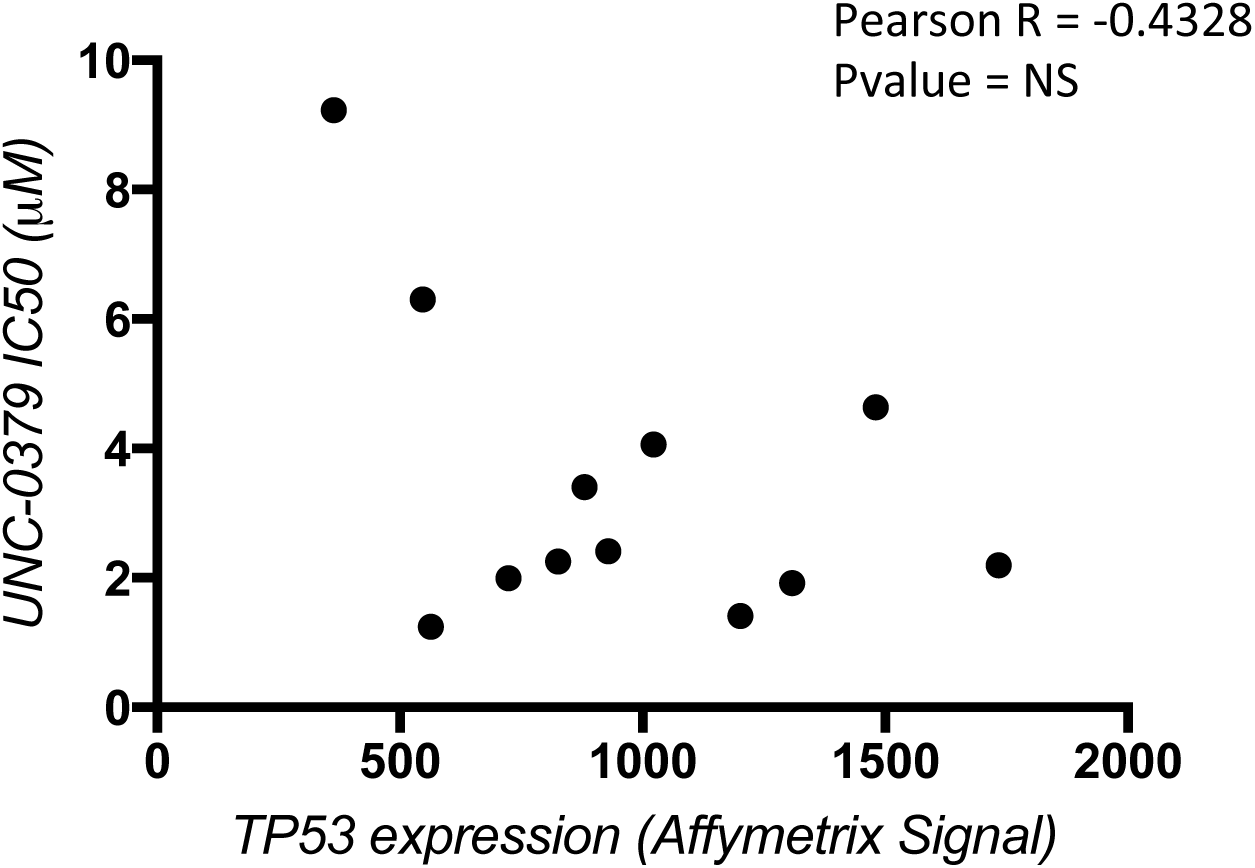
Correlation between *TP53* expression and drug response to UNC-0379 in HMCLs. IC50 was determined using CTG-based growth assay, *TP53* expression was obtained from Affymetrix microarrays data previously published.

**Supplementary Figure S9:**
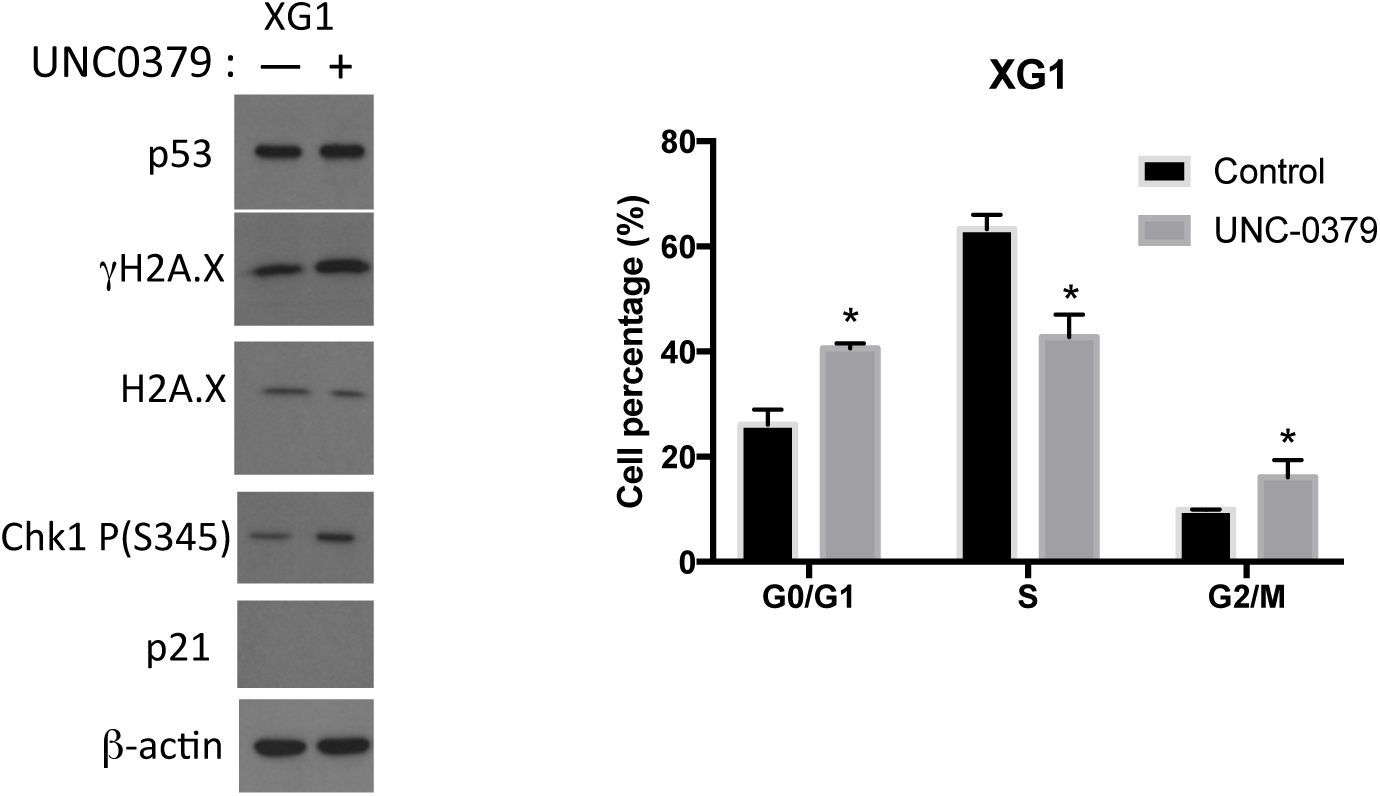
UNC-0379 induces genomic instability in XG1 p53-mutant cell line. **(A)** Immunoblot analysis of indicated proteins in total lysates from UNC-0379-treated (5μM) or untreated XG1 HMCL. β-actin and H2A.X were used as loading controls. **(B)** Cell cycle of control and 48h UNC-0379-treated (5μM) XG1 HMCL was analyzed by flow cytometry using DAPI, BrdU incorporation and labelling with an anti-BrdU antibody. Results are representative of three independent experiments. * indicates a significant difference compared to control cells using a Wilcoxon test for pairs (P ≤ 0.05).

**Supplementary Figure S10:**
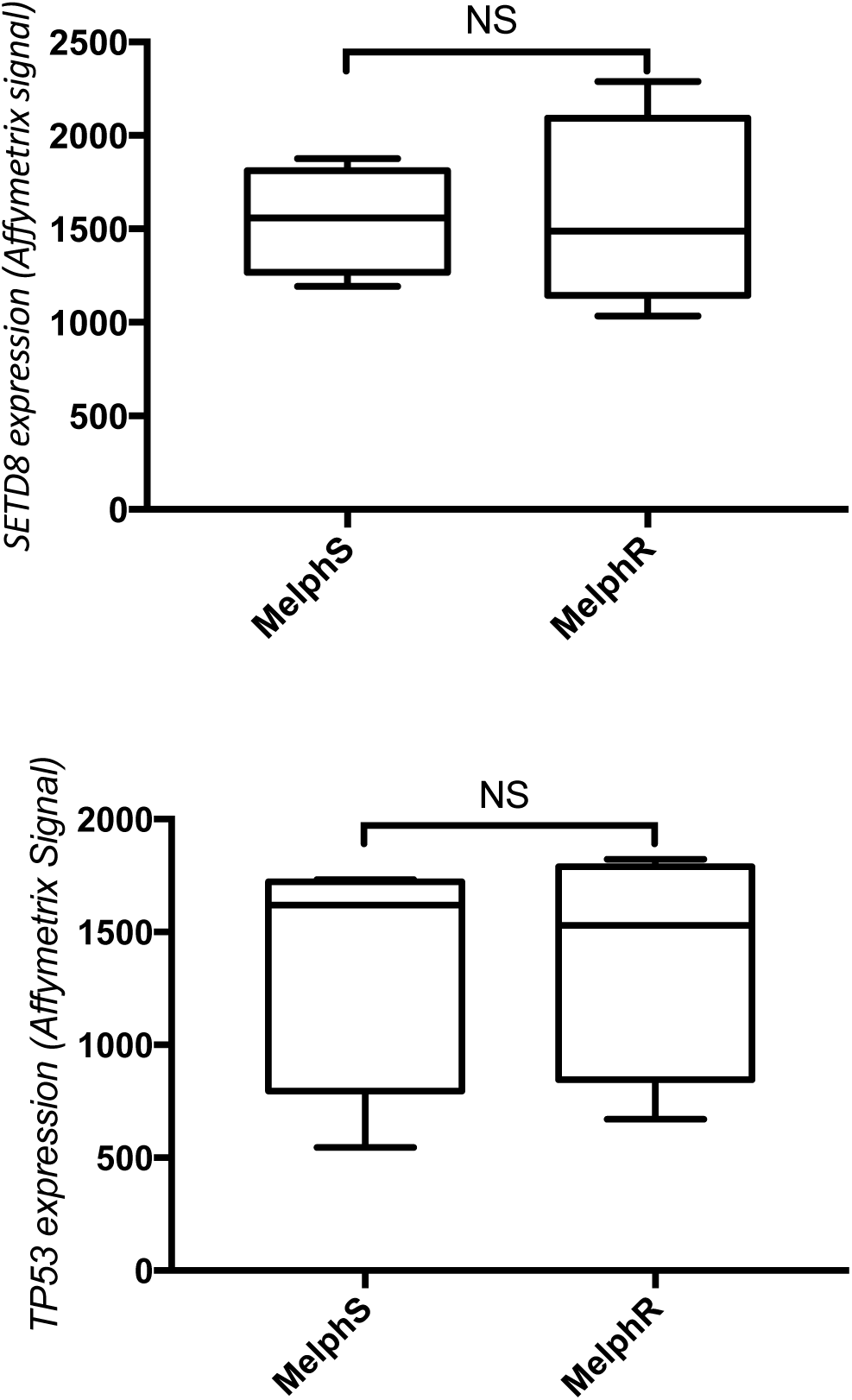
UNC-0379 induces genomic instability in XG1 p53-mutant cell line. Comparison of SETD8 and TP53 expression in XG7 HMCL sensitive (MelphS) or resistant (MelphR) to Melphalan treatment.

## ACKNOWLEDGMENTS

This work was supported by grants from labex Epigenmed, SIRIC-Montpellier-Cancer; Ligue Nationale Contre le Cancer (LNCC). Institutional Support was provided by the Institut National de la Santé et de la Recherche Médicale (INSERM) and by the Centre National de la Recherche Scientifique (CNRS). F.I. was supported by fellowships from LNCC and fondation pour la recherche médicale (FRM).

## Author contributions

EJ and JM designed the project, supervised the research, and wrote the paper. LH, FI, CG designed, performed the research and analyzed the data. LH and CG contributed to paper writing. OKC, CG, CB participated to the research LV and GC participated to clinical data analysis. AM and JJ contributed to design and production of SETD8 inhibitor. ED and EDB contributed to experiments with primary murine models.

